# Ophiuchus-Ab: A Versatile Generative Foundation Model for Advanced Antibody-Based Immunotherapy

**DOI:** 10.64898/2026.02.02.703197

**Authors:** Yiheng Zhu, Jian Ma, Mingze Yin, Jialu Wu, Lin Tang, Zhiyun Zhang, Qiuyi Li, Shikun Feng, Haiguang Liu, Tao Qin, Junchi Yan, Chang-Yu Hsieh, Tingjun Hou

**Author notes:** These authors contributed equally to this work (Yiheng Zhu, Jian Ma, Mingze Yin, Jialu Wu).

## Abstract

Antibodies exhibit extraordinary specificity and diversity in antigen recognition and have become a central class of therapeutics across a wide range of diseases. Despite this clinical success, antibody design remains fundamentally challenging. Antibody function emerges from intricate and highly coupled interactions between heavy and light chains, which complicate sequence-function relationships and limit the rational design of developable antibodies. Here, we reveal that modeling antibody sequence space at the level of paired heavy and light chains is essential to faithfully capture inter-chain dependencies, enabling a deeper understanding of antibody function and facilitating antibody discovery. We present Ophiuchus-Ab, a generative foundation model pre-trained on largescale paired antibody repertoires within a diffusion language modeling framework, unifying antibody generation and representation learning in a single probabilistic formulation. This framework excels diverse antibody design tasks, including CDR infilling, antibody humanization, and light-chain pairing. Beyond generation, diffusion-based pre-training yields transferable representations that enable accurate prediction of antibody properties, including developability, binding affinity, and specificity, even in low-data regimes. Together, these results establish Ophiuchus-Ab as a versatile foundation model for modeling antibodies, providing a foundation for next-generation antibody-based immunotherapy.

## 1 Introduction

Antibodies are large, Y-shaped proteins produced by B cells that play a central role in the adaptive immune system by specifically recognizing and neutralizing foreign antigens, such as pathogens or malignant cells [1]. In most mammals, antibodies comprise paired heavy and light chains with variable and constant domains, and a flexible hinge region connecting the heavy chains [2]. The variable domains of the heavy and light chains associate to form the Fv fragment, within which three complementarity-determining regions (CDRs) on each chain collectively define the antigen-binding site, enabling highly specific and high-affinity recognition of diverse epitopes.

Owing to these properties, antibodies have become an indispensable class of therapeutics in modern immunotherapy [3–6]. Antibodies can directly target tumorassociated antigens [7], modulate immune checkpoints [8], or deliver cytotoxic payloads with high specificity [9]. Clinically approved examples include trastuzumab [10] for HER2-positive breast cancer and immune checkpoint inhibitors such as pembrolizumab [11] and nivolumab [12], which enhance T-cell-mediated antitumor responses by blocking PD-1 signaling. Beyond conventional monoclonal antibodies, bispecific antibodies [13] and antibody–drug conjugates [14] have further expanded therapeutic modalities by engaging multiple immune mechanisms or improving targeted cytotoxic delivery. These advances have positioned antibodies at the forefront of treatments for cancer, infectious diseases, and autoimmune disorders.

Despite their success, the discovery and optimization of therapeutic antibodies remain a complex puzzle. Traditional antibody discovery pipelines rely primarily on *in vivo* animal immunization or *in vitro* display technologies such as phage display [15, 16]. Although effective, these approaches are time-consuming, labor-intensive, and costly, often requiring extensive experimental iteration. Moreover, antibody development poses unique challenges beyond those encountered in general protein engineering, including long production cycles, batch-to-batch variability, and the need for humanization to reduce immunogenicity [17, 18]. These constraints limit the speed and scalability of antibody discovery and complicate the rational optimization of affinity, specificity, and developability.

Recent advances in artificial intelligence (AI) have begun to reshape antibody development by enabling data-driven modeling of antibody sequences, structures, and functions [19]. In parallel, rapid progress in experimental techniques—including cryogenic electron microscopy, deep mutational scanning, and single-cell sequencing—has produced unprecedented volumes of high-quality antibody data [20, 21]. The concurrent maturation of algorithms and large-scale experimental datasets creates a timely opportunity to apply deep learning for uncovering the principles governing antibody recognition and for complementing or reinforcing traditional experimental discovery pipelines.

Inspired by the impressive success of large language models (LLMs) [22, 23] in natural language processing [24, 25], the conceptual analogy between protein sequences and natural language has motivated the development of protein language models (PLMs) for sequence modeling [26, 27]. Most existing PLMs adopt one of two pre-training paradigms: bidirectional masked language modeling or autoregressive causal language modeling, exemplified by BERT-and GPT-based architectures [24, 25]. Representative BERT-based models, including ProtBert [28] and the ESM family [29, 30], learn rich contextual representations that support downstream tasks such as fitness, function, and structure prediction [29–31]. Complementarily, GPT-based models, such as Prot-GPT2 [32], ProGen2 [33], and xTrimoPGLM [34], excel at generating coherent and biochemically plausible protein sequences for design applications [35, 36]. Building on these advances, antibody-specific PLMs have emerged to capture the distinctive statistical and functional properties of immunoglobulin sequences [37–40]. Representative models such as AbLang [38], AntiBERTa [39], and IgLM [40] are trained either from scratch on antibody repertoires or by fine-tuning general PLMs on antibody-specific data, enabling the modeling of sequence patterns shaped by antibody diversification processes, including V(D)J recombination and somatic hypermutation. By incorporating functional and mutational characteristics largely absent from general protein corpora, antibody-specific pre-training yields representations that are more appropriate for immunoglobulins. Consistent with this, antibody-specific PLMs achieve improved performance across diverse antibody-centered tasks [40–42], underscoring the importance of antibody-specific pre-training for data-driven antibody engineering.

Although antibody-specific PLMs have shown promising results, fully exploiting their potential for antibody modeling remains challenging. First, most existing models are pre-trained on single antibody chains, treating heavy and light chains independently and therefore failing to capture their native heterodimeric organization and co-evolutionary dependencies [33, 37, 38, 40]. Although recent studies have begun to explore modeling natively paired antibody sequences [43–45], these approaches introduce only limited network architectural and modeling adaptations, leading to relatively modest improvements. Second, prevailing BERT-like and GPT-like pretraining paradigms are not well aligned with the biological characteristics of antibodies. Because antibody hypervariable regions are not evolutionarily conserved, BERT-style models that emphasize evolutionary context are suboptimal for capturing antibody-specific sequence variability [30]. Meanwhile, GPT-style autoregressive models are inherently ill-suited for sequence infilling tasks, such as CDR redesign, which are central to antibody engineering [33]. Finally, existing approaches typically emphasize either antibody sequence understanding [37, 38] or generation [40], but rarely unify both capabilities within a single framework, limiting their generality across diverse antibody discovery applications.

To bridge the gap, we propose Ophiuchus-Ab (Fig. 1), a generative antibody foundation model pre-trained on large-scale paired antibody repertoires that unifies antibody generation and representation learning within a single probabilistic framework. Ophiuchus-Ab formulates antibody sequence space at the level of paired heavy and light chains, explicitly modeling their coupled evolution and inter-chain dependencies. Built upon diffusion language modeling [46, 47], Ophiuchus-Ab simultaneously learns joint, marginal, and conditional distributions over antibody sequences, enabling flexible generation and editing under arbitrary conditioning. The bidirectional, noncausal nature of diffusion modeling is particularly well suited to antibody design, where functionally critical regions are distributed across the sequence and governed by global constraints. Beyond generation, diffusion-based pre-training endows Ophiuchus-Ab with transferable representations that support accurate prediction of antibody properties, even in low-data regimes.

**Fig. 1:**
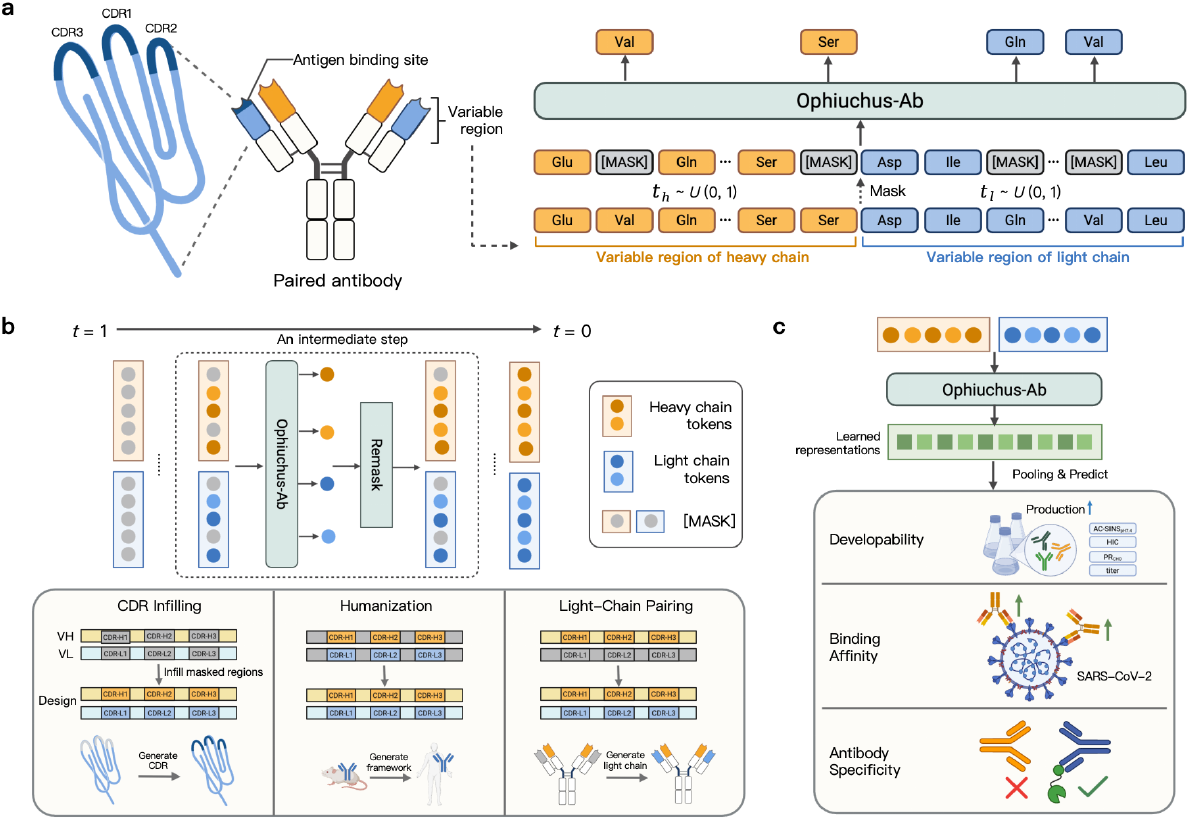
Overview of Ophiuchus-Ab, a versatile foundation model for unified antibody generation and understanding. (a) Ophiuchus-Ab is pre-trained on large-scale paired antibody repertoires within the generative probabilistic formulation of masked diffusion models. (b) Ophiuchus-Ab performs sampling by iteratively denoising masked regions of the paired sequence. By masking different regions, the same generative process supports multiple antibody design tasks, including CDR infilling, humanization, and light-chain pairing. (c) Beyond generation, diffusion-based pre-training yields transferable representations of paired antibody sequences, which are leveraged for downstream property prediction tasks, including developability, binding affinity, and binding specificity.

We systematically evaluate Ophiuchus-Ab across a broad range of antibody generation and understanding tasks, with results summarized in Fig. 2. Across antibody design applications, Ophiuchus-Ab consistently exhibits strong performance in CDR infilling, humanization, and light-chain pairing, demonstrating its ability to flexibly generate and edit antibody sequences under diverse constraints. In antibody understanding tasks, the representations learned by Ophiuchus-Ab enable accurate prediction of key antibody properties, including developability, binding affinity, and specificity. Notably, Ophiuchus-Ab consistently outperforms both general PLMs and existing antibody-specific PLMs across these tasks, highlighting the advantages of jointly modeling paired antibody chains within a unified diffusion-based framework. Together, these results establish Ophiuchus-Ab as a versatile antibody foundation model that advances both antibody design and understanding, providing a principled and effective framework for next-generation antibody-based immunotherapy.

**Fig. 2:**
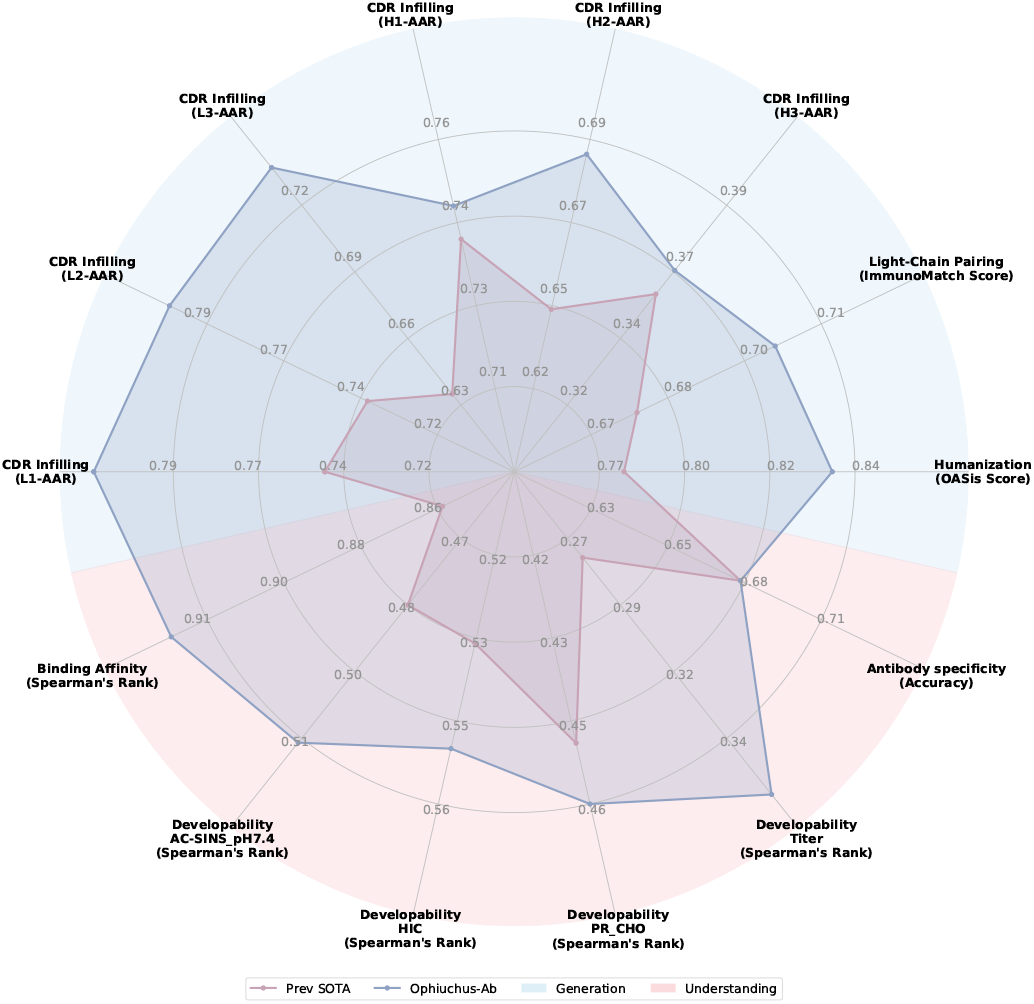
Ophiuchus-Ab outperforms previous state-of-the-art models across a wide range of generation and understanding tasks.

## 2 Results

### 2.1 Ophiuchus-Ab as a versatile antibody foundation model

Ophiuchus-Ab is developed as a versatile antibody foundation model, as illustrated in Fig. 1, through self-supervised learning on around 2.5 million paired antibody sequences from the Observed Antibody Space (OAS) database [21] (see section 4.1), enabling effective adaptation to a wide range of downstream understanding and generation tasks that are critical for antibody design. To achieve this purpose, we pre-train Ophiuchus-Ab within the generative probabilistic formulation of diffusion language models [46, 47]. After pre-training, Ophiuchus-Ab exhibits the ability to generate plausible, novel, and diverse antibody sequences in arbitrary orders, enabling a range of antibody design tasks, including CDR infilling, humanization, and light-chain pairing. Moreover, the diffusion-based generative pre-training endows Ophiuchus-Ab with a deeper understanding of antibody sequence patterns. Consequently, Ophiuchus-Ab can extract meaningful and generalizable representations for characterizing antibody functions and properties, even when labeled data are scarce, thereby providing a powerful tool for elucidating antibody mechanisms in a data-driven manner.

#### Paired antibody modeling with the inter-chain-aware backbone

A key design principle of Ophiuchus-Ab is to treat paired antibody sequences as the fundamental modeling unit. Because antibody function arises from tightly coupled interactions between heavy and light chains, modeling them independently obscures critical inter-chain dependencies. To capture these relationships, Ophiuchus-Ab is trained directly on natively paired antibody sequences and is equipped with a cross-chain attention mechanism built upon the pre-trained ESM-2 650M [30, 48]. This design preserves the rich intra-chain representations learned through large-scale protein pre-training while enabling explicit information exchange between chains. By jointly modeling intra- and inter-chain dependencies, Ophiuchus-Ab provides a more faithful representation of antibodies than single-chain formulations. Further architectural details are provided in section 4.2.1.

#### Diffusion language modeling as a unified probabilistic framework

Another key design principle of Ophiuchus-Ab is the adoption of a diffusion-based language modeling framework [46, 47] for antibody sequence modeling (see section 4.2.2). Unlike conventional BERT-like or GPT-like paradigms, diffusion language models define a flexible generative process that enables modeling of antibody sequences in arbitrary orders. In this formulation, Ophiuchus-Ab is cast as a masked token predictor for both heavy- and light-chain tokens, parametrized as *p*_*θ*_(*·*|*x*_*t*_), which takes a partially masked sequence *x*_*t*_ as input and predicts all masked tokens simultaneously. The model is trained using a cross-entropy loss computed only on the masked tokens:

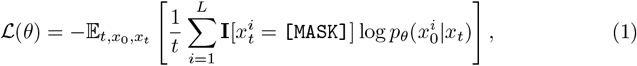

where *x*_0_ is ground truth, the timestep *t* is sampled uniformly from [0, 1], and *x*_*t*_ is obtained by masking *x*_0_. **I**[*·*] denotes the indicator function to ensure that the loss is computed only over the masked tokens.

To further improve modeling flexibility, learning efficiency, and robustness, we introduce three complementary design choices within the diffusion language modeling framework. First, we assign independent noise levels to the heavy and light chains, enabling flexible sampling across diverse antibody design tasks (section 4.2.3). Second, to emphasize functionally relevant, non-germline residues [49], we replace the standard cross-entropy objective used in diffusion language models with focal loss [50], which down-weights well-predicted tokens (section 4.2.4). Third, we adopt complementary masking [51] to improve data efficiency by ensuring that each token position contributes to training, thereby enhancing sample efficiency and optimization stability (section 4.2.5).

#### Unified generation and understanding for generalist antibody design

Enabled by diffusion-based probabilistic learning and strengthened by inter-chain modeling, Ophiuchus-Ab unifies antibody sequence generation and representation learning within a single foundation model. As a generative model, Ophiuchus-Ab samples novel antibody sequences using a diffusion-based process (section 4.2.6) rather than a left-to-right autoregressive scheme. Generation is achieved by reversing the diffusion process from fully or partially masked inputs (Fig. 1b). At intermediate steps, Ophiuchus-Ab simultaneously predicts all masked tokens and preferentially re-masks low-confidence predictions until all positions are resolved [52]. This inference flexibility enables both unconditional sequence generation and targeted infilling of regions such as CDRs, supporting a wide range of antibody design tasks without requiring task-specific sampling strategies. As a representation model, Ophiuchus-Ab maps paired antibody sequences to informative embeddings that capture sequence-level characteristics (Fig. 1c). Given a paired antibody sequence as input, the pre-trained model produces token-level representations that are mean-pooled to yield a fixed-length vector. These embeddings encode both local sequence features and global paired antibody context, providing a compact and transferable representation for downstream prediction tasks.

### 2.2 Ophiuchus-Ab generates natural, novel, and diverse antibodies *in silico*

Our proposed Ophiuchus-Ab, built upon diffusion language models, provides more flexible inference than autoregressive PLMs in two key aspects. First, Ophiuchus-Ab is not limited to left-to-right generation and can generate outputs in arbitrary orders, enabling a wider range of antibody design tasks, from sequence infilling to *de novo* design. Second, the number of tokens generated at each step can be adjusted dynamically, offering a tunable trade-off between computational efficiency and sample quality. This flexibility constitutes distinct advantages over traditional autoregressive frameworks.

In this section, we demonstrate the generative capabilities of Ophiuchus-Ab across several practically meaningful antibody design scenarios, including CDR infilling, humanization, and light-chain pairing.

#### 2.2.1 CDR infilling

Each antibody chain contains three CDRs, which play a critical role in determining binding affinity to a target antigen, while having relatively minor effects on other structural properties of the antibody. Thus far, antibody design methods have focused on redesigning the CDRs while leaving the rest of the antibody fixed, essentially treating it as an infilling task [40, 53, 54]. Redesigning the CDRs is a crucial aspect of antibody development. This process not only optimizes existing antibodies to improve their binding affinity and specificity [53], but also facilitates the creation of diverse loop libraries [40]. Such strategy holds substantial promise for advancing antibody-based therapeutics with enhanced precision.

The any-order nature of generation enables Ophiuchus-Ab to be applied in a zero-shot manner to the CDR infilling task, despite not being specifically trained for this conditional generation task. Specifically, we first mask the CDR regions and then decode the masked regions of the sequence, without requiring fine-tuning or modifications to the sampling strategy.

The ability of Ophiuchus-Ab to recover CDRs is evaluated on the Structural Antibody Database (SAbDab) [20]. We consider two test sets derived from the SAbDab in our analysis: (i) the SAb23H2 [55] which includes 60 antigen-antibody complexes determined experimentally and published between June 30, 2023, and December 30, 2023, respectively. (ii) The SAbDab dataset collected in Kong et al. [56], which contains 3,127 antigen-antibody complexes after filtering structures that lack light chains or antigens. To prevent data leakage, our pre-training dataset is filtered to remove sequences similar to those in either test set. For dataset details, see section 4.1. We evaluate performance using Amino Acid Recovery (AAR), defined as the fraction of correctly recovered amino acids in the infilled regions relative to the ground-truth sequence.

As shown in table 1 and table 2, Ophiuchus-Ab exhibits strong zero-shot capability for the CDR infilling task. On the SAb23H2 dataset, the model achieves the highest AAR across both light- and heavy-chain CDRs, outperforming existing approaches such as dyMEAN [57] and IgGM [55]. Notably, whereas several competing methods rely on antibody structures predicted by external models (e.g., IgFold [58] or AlphaFold3 [59]), Ophiuchus-Ab operates purely at the sequence level and nonetheless attains superior performance, highlighting the effectiveness of paired heavy–light chain pre-training. On the SAbDab benchmark, which evaluates CDR infilling exclusively on the heavy chain, Ophiuchus-Ab achieves state-of-the-art performance on CDR-H3 and yields competitive results on CDR-H1 and CDR-H2. Although BERT-based PLMs such as AntiBERTy [37] and AbLang2 [49] report slightly higher AAR on CDR-H1 and CDR-H2, this advantage may arise from their orders-of-magnitude larger unpaired heavy-chain pre-training datasets, which particularly favor the less diverse CDR-H1 and CDR-H2 regions, as well as from potential overlap between their pre-training data and SAbDab.

**Table 1:**
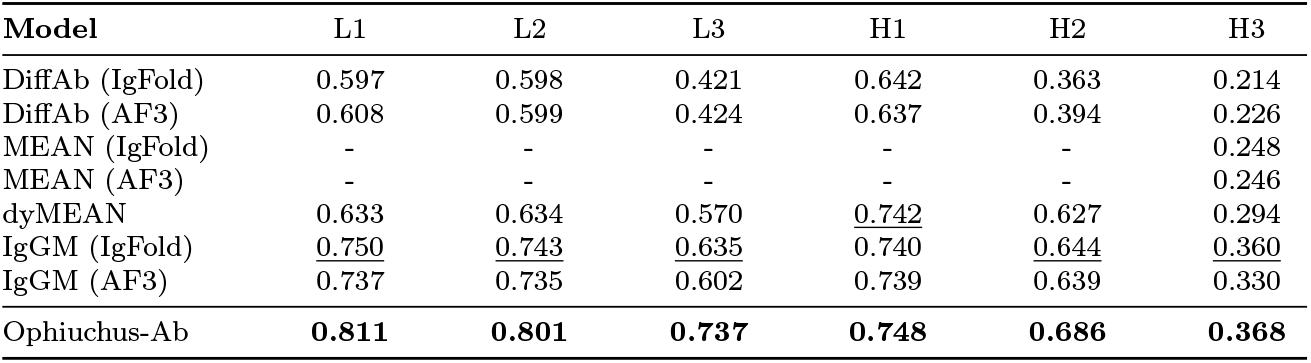
Results of CDR infilling on the SAb23H2 dataset. Methods annotated with (IgFold) or (AF3) use antibody structures predicted by IgFold or AlphaFold3, respectively, as the initial input. H1-H3 indicate the CDRs of heavy chain while L1-L3 indicate the CDRs of light chain. **Bold** and underline denote the best and second metrics.

**Table 2:**
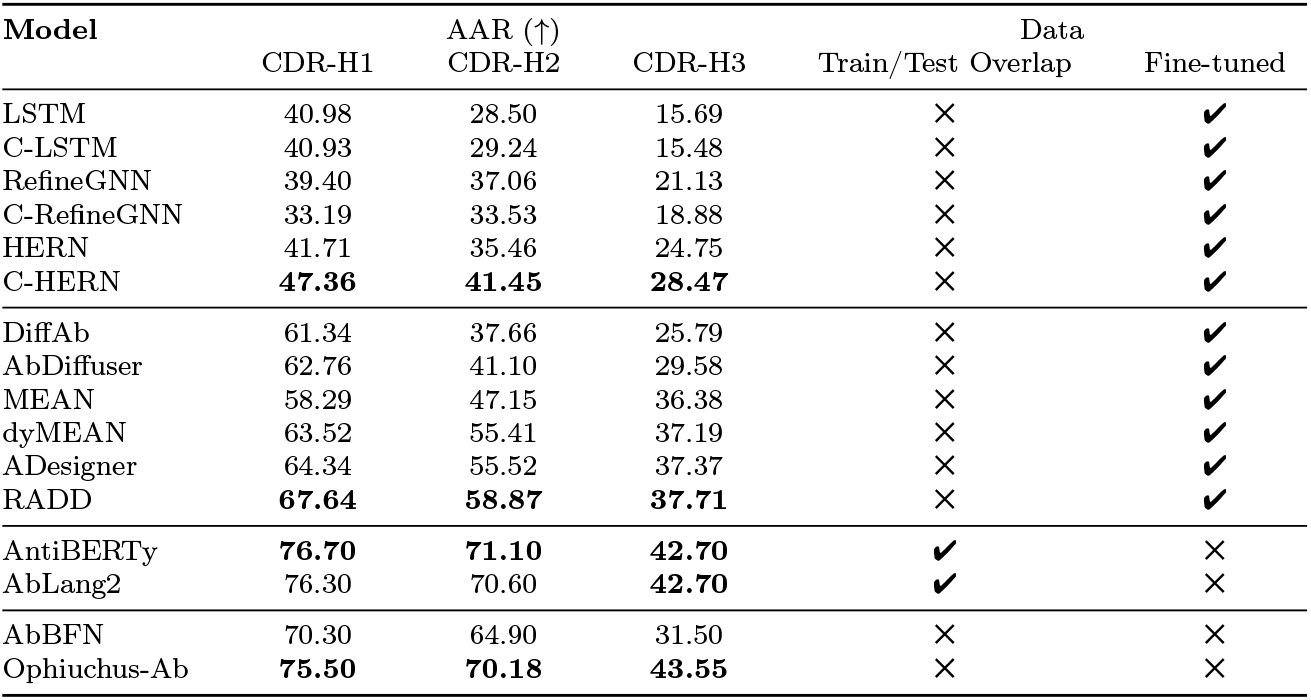
Mean CDR infilling performance on the SAbDab dataset evaluated using 10-fold cross-validation. “Train/Test Overlap” indicates whether a model’s pre-training data were filtered to prevent data leakage. “Fine-tuned” indicates whether the model was subsequently fine-tuned on the nine training folds corresponding to each test fold.

#### 2.2.2 Humanization

Non-human–derived antibodies can elicit anti-drug immune responses, making humanization a critical step in therapeutic antibody development [60]. Traditional antibody engineering strategies typically humanize non-human antibodies through framework residue substitution or CDR grafting onto human scaffolds [17, 18]. Although these approaches have enabled many successful therapeutics, they often require substantial expert knowledge and iterative experimental refinement to reduce immunogenicity while preserving affinity, specificity, and overall developability [61]. More recently, deep learning–based approaches have emerged as an alternative paradigm for antibody humanization. Representative examples include AbNatiV [59], Humatch [62], and CUMAb [63]. AbNatiV and Humatch formulate humanization primarily as a sequence-level residue substitution problem, whereas CUMAb explicitly incorporates structural constraints to account for three-dimensional context.

Building on these advances, we examine the humanization capability of Ophiuchus-Ab. Inspired by Ma et al. [64], we cast antibody humanization as a conditional infilling task in which the CDRs are fixed and the framework residues are generated by the model. To promote human-like sequence characteristics and avoid the introduction of non-human patterns, we fine-tune Ophiuchus-Ab exclusively on human antibody sequences. This training strategy enforces a human-specific prior over antibody sequence space, minimizing cross-species signal leakage and guiding the reconstruction of framework regions toward human-compatible solutions.

We benchmark the humanization performance of Ophiuchus-Ab against HuDiff [64] and IgCraft [65]. Following Greenig et al. [65], we evaluate all methods on a test set of 27 murine antibodies from SAbDab [20]. Given IMGT-annotated CDRs as fixed inputs, Ophiuchus-Ab further generates the corresponding framework regions. Humanization quality is assessed using the OASis humanness score [66] and the AbNatiV score [67], with results summarized in Fig. 3. Across all test antibodies, Ophiuchus-Ab achieves substantially higher humanness than both baselines, increasing the OASis score from 47.3 for the original murine sequences to 83.4, compared with 74.6 for HuDiff and 77.9 for IgCraft. Consistent trends are observed with AbNatiV score, where Ophiuchus-Ab attains state-of-the-art scores on both chains. This indicates that, under identical CDR constraints, the frameworks generated by Ophiuchus-Ab are consistently closer to human antibody repertoires. In addition, Ophiuchus-Ab exhibits lower sequence identity to the original murine antibodies than baseline methods (approximately 65% versus 80%), reflecting broader framework substitutions that accompany the observed gains in humanness. Overall, Ophiuchus-Ab produces highly human-like framework design while introducing a controlled and biologically plausible degree of sequence divergence under fixed CDR constraints.

**Fig. 3:**
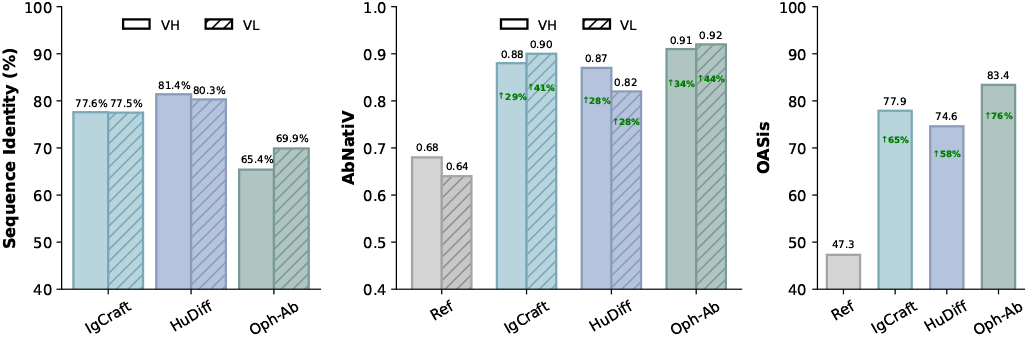
Humanization benchmarking on 27 murine antibodies from SAbDab comparing IgCraft, HuDiff, and Ophiuchus-Ab (Oph-Ab), with VH/VL sequence identity to the parental murine sequences, AbNatiV humanness scores for each chain, and OASis humanness scores for the full antibody sequence.

#### 2.2.3 Light-chain pairing

The adaptive immune system generates a highly diverse antibody repertoire to recognize a wide range of antigens. This diversity arises from V(D)J recombination and somatic mutation in both heavy and light chains. Although antibodies are often treated as modular assemblies, the specificity of heavy–light pairing remains incompletely understood [68]. Accumulating evidence shows that productive antibody function depends on coordinated interactions between the CDRs and framework regions of both chains, which jointly shape antigen recognition and binding stability [69]. At the same time, immunological studies indicate that during B-cell maturation, a given heavy chain can transiently pair with multiple light chains before a functional combination is selected [2]. Together, these observations reveal a fundamental tension between pairing flexibility and functional constraint, underscoring the importance of accurately identifying cognate light-chain partners in therapeutic antibody design. Accordingly, computational approaches have been developed to address this challenge either by predicting compatible heavy–light pairs or by generating complementary light chains from its pairing partner [44, 68, 70].

To assess whether Ophiuchus-Ab captures heavy–light pairing constraints, we evaluate its performance on the light-chain pairing task, in which the model designs cognate light-chain partners conditioned on a given heavy chain. We benchmark Ophiuchus-Ab against p-IgGen [44] and LiChen [70] using a comprehensive evaluation protocol. Pairwise complementarity between generated light chains and their conditioned heavy chains is quantified using ImmunoMatch scores [68], and we report the proportion of cases in which the generated pair exceeds the corresponding reference pair. We further assess biological consistency by evaluating agreement with the reference chain type (*λ* versus *κ*) and V/J gene assignments at both the gene and family levels. Antibody likeness is assessed using statistical metrics that capture sequence diversity and validity, defined by the successful annotation of sequences by ANARCI [71], as well as the normalized average Wasserstein distance W_property_ between generated and reference distributions of antibody-related properties, including hydrophobicity, charge, and sequence length, among other properties derived from the sequence with Biopython [72, 73].

We evaluate all methods on a held-out test set comprising 500 sequences. For each heavy chain, eight candidate light-chain sequences are generated by each model, and the results are summarized in table 3. We begin by considering the setting in which light-chain sequences are generated conditioned solely on the heavy chain. Ophiuchus-Ab consistently outperforms p-IgGen and LiChen in ImmunoMatch score, indicating stronger heavy–light compatibility. Agreement with the reference light chains, however, remains low across all methods, consistent with the intrinsic diversity of antibody pairing, whereby a single heavy chain can associate with multiple functionally compatible light-chain partners. These results demonstrate that Ophiuchus-Ab can generate a diverse set of plausible light-chain partners for a given heavy chain, capturing the inherent flexibility of heavy–light pairing in the absence of light-chain constraints. We further evaluate a partially conditioned setting in which light-chain generation is guided by the heavy chain together with the first three residues of the light chain. This minimal prompting substantially increases agreement with the reference across all methods, with the most pronounced gains observed in V-family matching, indicating that short N-terminal cues effectively constrain chain-type and germline usage. Under this setting, Ophiuchus-Ab remains consistently stronger than the baselines and continues to achieve the highest ImmunoMatch score.

**Table 3:** Light-chain pairing performance with and without initial three-residue prompting. Ophiuchus-Ab is benchmarked against p-IgGen [44] and LiChen [70] using metrics that assess pairing compatibility, biological consistency, and antibody likeness.

| Method | Pairing Score |  | Biological Consistency |  |  |  |  | Antibody likeness |  |  |
| --- | --- | --- | --- | --- | --- | --- | --- | --- | --- | --- |
|  | ImmunoMatch (↑) | Better (↑) | Chain | V | J | V-fam | J-fam | Div. (↑) | Valid. (↑) | W<sub>property</sub> (↓) |
| **Without initial-residue prompting** |  |  |  |  |  |  |  |  |  |  |
| LiChen | 0.657 | 46.2% | 62.3% | 0.049 | 0.156 | 0.215 | 0.157 | 0.515 | **100.0%** | 0.140 |
| p-IgGen | 0.674 | 50.5% | 50.5% | 0.045 | 0.139 | 0.171 | 0.142 | **0.709** | 99.1% | 0.103 |
| Ophiuchus-Ab | **0.701** | **51.6%** | 55.6% | 0.056 | 0.150 | 0.169 | 0.152 | 0.676 | **100.0%** | **0.072** |
| **With initial 3-residue prompting** |  |  |  |  |  |  |  |  |  |  |
| LiChen | 0.633 | 42.7% | **99.7%** | 0.298 | **0.290** | 0.890 | **0.291** | 0.254 | 99.9% | 0.150 |
| p-IgGen | 0.669 | 45.3% | 98.4% | 0.313 | 0.283 | 0.874 | 0.288 | **0.390** | 99.2% | 0.118 |
| Ophiuchus-Ab | **0.695** | **49.2%** | **99.7%** | **0.356** | 0.273 | **0.894** | 0.276 | 0.335 | **100.0%** | **0.066** |

Comparing the unconditioned and partially conditioned settings reveals a trade-off. While first three-residue prompting improves agreement with the reference, it leads to a modest reduction in ImmunoMatch score, indicating that additional conditioning narrows the space of compatible light chains toward the recorded reference sequence. Importantly, these results indicate that the reference light chain represents only one of multiple biologically plausible partners for a given heavy chain. By capturing this intrinsic flexibility, Ophiuchus-Ab is well suited for practical antibody discovery, enabling the exploration of alternative compatible light chains and the systematic expansion of paired antibody repertoires beyond those observed in existing datasets.

### 2.3 Ophiuchus-Ab learns meaningful and generalizable representations for understanding antibody

Recently, PLMs pre-trained on natural protein sequences have been remarkably successful at learning rich, context-dependent representations of amino acid sequences, enabling a deeper understanding of the proteins [30]. These learned representations have been shown to capture evolutionary relationships and support a wide range of downstream tasks, from function annotation [29] and structure prediction [30] to protein engineering [74].

In this section, we evaluate whether Ophiuchus-Ab can extract meaningful and generalizable representations for characterizing antibody functions and properties. We systematically assess the model’s performance on a series of property prediction tasks that are critical to antibody design, including developability [75], binding affinity [76], and specificity [77]. This structured evaluation, progressing from fundamental to increasingly complex properties, is intended to comprehensively determine the model’s ability to capture essential aspects of antibody functionality and overall viability.

#### 2.3.1 Developability

As development progresses from clinical trials to commercialization, the antibody developability significantly influences the speed, cost, and likelihood of successful commercialization [78]. Consequently, conducting developability assessments early in the drug discovery process is vital for risk mitigation [75, 79]. Antibody developability encompasses both *in vitro* properties, such as high-titer expression in heterologous systems, high solubility, thermostability, and minimal self-association and viscosity, as well as *in vivo* characteristics, including low immunogenicity, reduced non-specific “sticky” binding, and a prolonged half-life [80, 81]. These attributes are crucial for ensuring manufacturability and clinical suitability [82].

Experimental assessment of developability profiles is both costly and time-consuming. Traditional computational tools, such as the Therapeutic Antibody Profiler (TAP) [75], provide five descriptors for antibody variable regions, capturing features such as the total length of the CDRs, the surface hydrophobicity, positive charge and negative charge in the CDRs, and asymmetry in the net heavy- and light-chain surface charges. However, these hand-crafted features are insufficient for accurately predicting developability, which is why deep learning methods have become increasingly important.

To demonstrate that Ophiuchus-Ab can play a critical role in early-stage risk assessment of therapeutic antibody candidates, we conduct experiments on the GDPa1 benchmark [83] to evaluate whether the antibody representations learned by Ophiuchus-Ab enable accurate developability prediction. The GDPa1 dataset comprises paired heavy- and light-chain variable regions from 246 clinical-stage antibodies, along with biophysical assay measurements characterizing their developability profiles. We consider four key antibody developability properties: (1) AC-SINS at pH 7.4 (**AC-SINS**_**pH7.4**_), which measures self-interaction propensity; (2) hydrophobic interaction chromatography (**HIC**), a proxy for hydrophobicity and aggregation propensity; (3) polyreactivity in Chinese hamster ovary (CHO) cells (**PR**_**CHO**_), which reflects non-specific binding to off-target proteins; and (4) **titer**, representing antibody expression yield in mammalian cells.

The pre-trained Ophiuchus-Ab takes paired antibody sequences as input and outputs representations. For each developability property, a ridge regression model is trained on these representations. We employ 5-fold cross-validation with isotype-stratified folds, ensuring that evaluation is conducted on novel antibody families. Model performance is measured using Spearman’s rank correlation coefficient. We systematically compare Ophiuchus-Ab against three families of baselines. One-hot encoding, TAP [75], and DeepSP [84] are descriptor-based methods. ESM-2 [30] is a general PLM. p-IgGen [44] and AbLang-2 [49] are antibody-specific PLMs.

As presented in table 4, Ophiuchus-Ab consistently outperforms all baseline methods across the four tasks, achieving an average relative improvement of 11.9% over the previous state-of-the-art model, AbLang-2. We highlight the following observations. (1) Both general PLMs and antibody-specific PLMs substantially outperform descriptor-based methods, indicating that large-scale pre-training enables effective transfer to antibody-related tasks. In contrast, one-hot encoding and TAP perform poorly across all properties. DeepSP, which introduces spatial features derived from antibody sequences, shows some advantage in predicting HIC but still underperforms relative to PLMs overall. (2) Antibody-specific PLMs achieve superior overall performance compared with general PLMs, suggesting that focused pre-training on antibody sequences is critical for improving developability prediction. Across all four properties, in addition to Ophiuchus-Ab, AbLang-2 also demonstrates promising performance.

**Table 4:** Mean results of 5-fold cross-validation with isotype-stratified folds on the GDPa1 dataset, where bold and underline denote the best and second metrics.

| Model | AC-SINS_pH7.4 | HIC | PR_CHO | Titer |
| --- | --- | --- | --- | --- |
| One-hot encoding | 0.230 | 0.233 | 0.204 | 0.193 |
| TAP | 0.294 | 0.222 | 0.136 | 0.113 |
| DeepSP | 0.348 | <u>0.531</u> | 0.257 | 0.114 |
| ESM-2 | 0.420 | 0.416 | 0.420 | 0.180 |
| p-IgGen | 0.388 | 0.346 | 0.424 | 0.238 |
| AbLang-2 | <u>0.480</u> | 0.472 | <u>0.449</u> | <u>0.279</u> |
| Ophiuchus-Ab | **0.511** | **0.550** | **0.460** | **0.359** |

#### 2.3.2 Binding affinity

Beyond antibody developability, we evaluate the performance of Ophiuchus-Ab in predicting binding affinity, an immune-related property that is critical for therapeutic development. Binding affinity prediction is a central task in computational antibody design and optimization, particularly in low-N settings, where the objective is to extrapolate the effects of combinatorial mutations from a limited set of assay-labeled antibodies to a broad space of candidate variants [85]. Accurate and reliable affinity predictions enable the prioritization of promising mutations and guide subsequent experimental assays [86].

We apply Ophiuchus-Ab to predict changes in binding affinity for m396 antibody mutants against SARS-CoV-2, using experimental data from Desautels et al. [87]. The wild-type m396 antibody targets the receptor binding domain (RBD) of the SARS-CoV-1 spike protein [88], and extensive in silico mutagenesis was performed to assess its potential binding to the SARS-CoV-2 RBD. In total, approximately 90,000 mutants were evaluated, with binding affinities estimated via ΔΔ*G* scores computed using five established energy functions: FoldX Whole, FoldX Interface Only [89], Statium [90], Rosetta Flex, and Rosetta Total Energy [91].

We compare Ophiuchus-Ab with AbMAP [41], a transfer learning framework that adapts general PLMs to antibody-specific tasks, and with MINT [48], a PLM specifically designed to model protein–protein interactions. To align with the low-N setting characteristic of real-world antibody design, we adopt the dataset splitting strategy used by AbMAP. This involves training on a small subset of the data and evaluating performance on the remaining portions, with results measured using Spearman’s rank correlation coefficient. As shown in Fig. 4, Ophiuchus-Ab outperforms AbMAP and MINT across all data splits, with the improvement becoming increasingly pronounced under limited-data conditions. Remarkably, when trained on only 0.5% of the data, Ophiuchus-Ab exceeds the previous state-of-the-art model, MINT, by over 6%. This result highlights the strong potential of Ophiuchus-Ab for effective antibody design in low-data regimes.

**Fig. 4:**
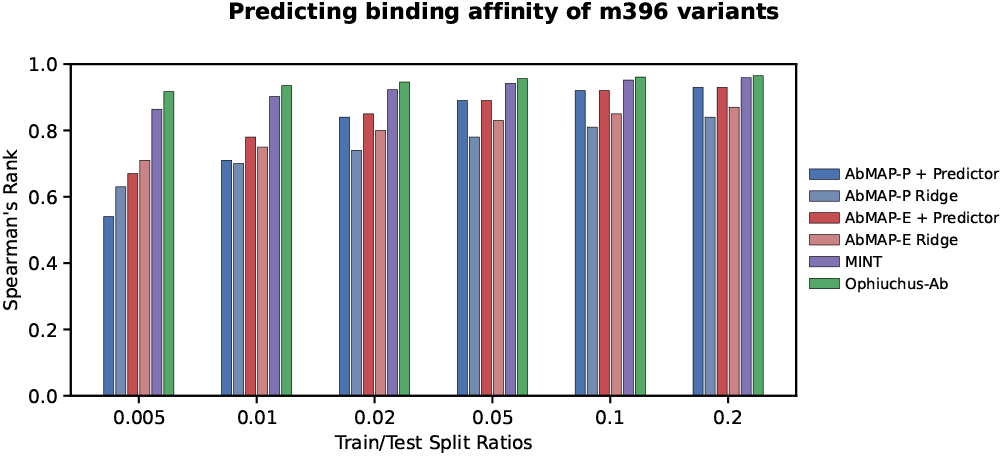
Comparative evaluation of protein representations for binding affinity prediction in a low-N context. AbMAP-P refers to AbMAP with the ProtBert model and AbMAP-E refers to the ESM-1b version. The two AbMAP models are also evaluated using MLP and ridge regression models for prediction.

#### 2.3.3 Antibody specificity

Antibody specificity reflects the immune system’s ability to selectively recognize antigens while avoiding nonspecific or off-target interactions, and is therefore central to the identification and optimization of therapeutic antibodies targeting specific pathogens [92]. We evaluate this capability using a three-way classification task that distinguishes coronavirus (CoV)-targeting antibodies, influenza (Flu)-targeting antibodies, and nonspecific healthy donor (HD) antibodies [93]. In this setting, CoV- and Flu-specific antibodies represent pathogen-targeted responses, whereas HD antibodies serve as negative controls to assess specificity and exclude nonspecific binding.

We evaluate Ophiuchus-Ab using linear classification on frozen representations, training a lightweight classification head while keeping all pre-trained parameters fixed. This setup isolates the quality of the learned representations and enables a fair assessment of antibody specificity. The model is evaluated on 4,398 paired antibody sequences with equal numbers per class, and the dataset is split using 5-fold stratified cross-validation. Performance is measured using accuracy, F1 score, and Matthews correlation coefficient (MCC). We compare Ophiuchus-Ab against a range of antibody-specific PLMs, including IgBERT [45], IgT5 [45], AbLang2 [49], AntiBERTa2 [94], and CurrAb [93].

As shown in table 5, Ophiuchus-Ab achieves state-of-the-art performance on antibody specificity classification, outperforming most existing antibody language models across evaluation metrics. Notably, Ophiuchus-Ab attains performance comparable to the strongest baseline, CurrAb, despite substantial differences in training complexity. While CurrAb relies on large unpaired datasets, curriculum learning, and extensive hyperparameter optimization, Ophiuchus-Ab achieves competitive results without task-specific tuning, indicating that it learns robust and transferable representations for antibody specificity.

**Table 5:** Performance comparison for antibody specificity classification (HD vs Flu vs CoV). Models are evaluated using Accuracy, F1 score, and Matthews correlation coefficient (MCC). Bold and underline denote the best and second-best results.

| Model | Accuracy (↑) | F1 (↑) | MCC (↑) |
| --- | --- | --- | --- |
| IgBERT | 0.5266 | 0.5226 | 0.2926 |
| IgT5 | 0.4789 | 0.4534 | 0.2390 |
| AbLang2 | 0.5780 | 0.5777 | 0.3675 |
| AntiBERTa2 | 0.5759 | 0.5768 | 0.3658 |
| CurrAb | **0.6796** | **0.6801** | <u>0.5200</u> |
| Ophiuchus-Ab | **0.6796** | <u>0.6790</u> | **0.5203** |

## 3 Discussion

In this work, we present Ophiuchus-Ab, an antibody foundation model built on diffusion language modeling [46, 47] that formulates antibody sequence space at the level of paired heavy and light chains and unifies antibody generation and representation learning through a probabilistic denoising process. Ophiuchus-Ab supports diverse antibody design tasks, including CDR infilling, humanization, and light-chain pairing, while producing transferable representations for downstream property prediction in low-data regimes.

### Limitations

Despite these advances, several limitations remain. First, like most masked diffusion models, Ophiuchus-Ab has limited capacity for controlled variable-length generation [95], making it challenging to model the insertions and deletions that commonly arise during antibody evolution. Second, the standard masked diffusion formulation lacks an inherent self-correction mechanism, whereby decoded tokens remain fixed during inference, restricting the model’s ability to correct suboptimal predictions that may compromise residue compatibility, inter-chain interactions, or functional motifs in antibody sequences. Third, Ophiuchus-Ab is trained to solve an exponentially large space of infilling problems under diverse masking patterns, whereas inference typically proceeds via a single, heuristically chosen decoding order [52]. In the context of antibody design, determining task-adaptive generation orders that improve sample quality remains an open question.

### Future directions

There are several promising directions for future research. Incorporating structural information into Ophiuchus-Ab could support more comprehensive antibody understanding and generation [42], for example by augmenting the model with structural constraints for inverse folding [96–98]. Moreover, the generation process of Ophiuchus-Ab can be naturally viewed as a Markov decision process, enabling integration with reinforcement learning [99] to support goal-directed antibody design by explicitly optimizing objectives such as binding affinity, specificity, and developability [100]. Finally, extending Ophiuchus-Ab to antigen-specific antibody design would facilitate practical applications by directly conditioning generation on target antigens [101].

## 4 Methods

### 4.1 Dataset curation

We collected paired antibody variable-region sequences from the Observed Antibody Space (OAS) database [21], comprising heavy-chain (VH) and light-chain (VL) variable domains.

#### Sequence filtering

Records containing ambiguous or non-standard amino acid residues were removed, and the remaining sequences were deduplicated based on exact VH–VL pairs. All VH and VL sequences were numbered using ANARCI with the IMGT scheme. We then applied sequential quality-control filters, first requiring the presence of conserved cysteines at IMGT positions 23 and 104 in both VH and VL chains, and subsequently enforcing terminus completeness by allowing at most three missing residues within the N- and C-terminal framework regions (FR1 and FR4) of each chain. To prevent data leakage in downstream evaluations, all remaining sequences were compared against the Structural Antibody Database (SAbDab) using MMseqs2 [102], and any sequence with greater than 95% identity to a SAbDab entry was excluded. After filtering, the final pre-training dataset comprised 2,511,612 unique paired VH–VL sequences.

#### Sequence Clustering

To construct non-redundant data splits, we clustered the filtered sequences following the protocol used in AbLang2 [49] and AbBFN2 [103]. Briefly, CDR3 sequences were extracted and used to define initial clusters, with each unique CDR3 forming a cluster. Within each CDR3 cluster, full-length variable domains were further clustered using Linclust at a 95% sequence identity threshold with cov-mode 1 [104]. Heavy and light chains were then paired, and two antibodies were considered to belong to the same final cluster if they shared identical CDR-H3 and CDR-L3 loops and exhibited greater than 95% sequence identity in both VH and VL domains. Based on these final clusters, we constructed the training, validation, and hold-out test sets by sampling 99%, 0.5%, and 0.5% of the unique clusters, respectively.

### 4.2 Overview of Ophiuchus-Ab

#### 4.2.1 Model Architecture

Ophiuchus-Ab is a Transformer-based antibody language model that explicitly models paired heavy–light chains as a single unit, enabling the capture of inter-chain interaction patterns. We build on the pre-trained ESM-2 650M model [30], which provides strong single-chain protein representations and has been widely adopted in protein modeling.

Given a paired antibody consisting of a heavy chain 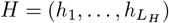 and a light chain 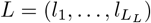, we tokenize the two chains independently using the standard vocabulary of ESM-2. We pad heavy and light chains to fixed maximum lengths *T*_*H*_ and *T*_*L*_, respectively (e.g., *T*_*H*_ = 150 for VH and *T*_*L*_ = 128 for VL), using special |EOS| tokens:

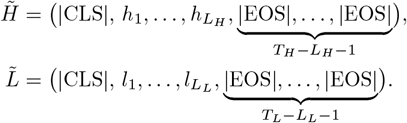

The final model input is formed by concatenating the two padded chains,

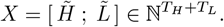

The |EOS| token is treated as a normal token during training and is removed during sampling, enabling Ophiuchus-Ab to automatically control chain length.

When paired antibody sequences are modeled by ESM-2 in a simple concatenated form, the model is applied outside its single-chain pre-training distribution. In this setting, residues from the heavy and light chains are handled by a shared self-attention mechanism, preventing the model from distinguishing inter-chain from intra-chain interactions and thereby entangling the two. Following Ullanat et al. [48], Liu et al. [105], we address this limitation by explicitly decoupling intra-chain and inter-chain attention. We introduce separate attention parameters for residue pairs within the same chain and across different chains, both initialized from the ESM-2 checkpoint and optimized independently during training. This design allows Ophiuchus-Ab to progressively capture heavy–light interaction patterns while retaining the strong single-chain representations inherited from ESM-2.

#### 4.2.2 Diffusion Language Modeling

Diffusion models define generative processes through a gradual noising of data followed by a learned denoising procedure that recovers samples from noise [106, 107]. For continuous variables, this is formalized as a forward Markov chain:

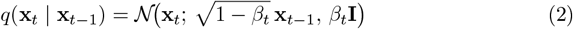

with a corresponding learned reverse process

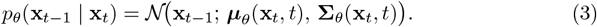

While originally developed for continuous domains such as images, a class of discrete variants, including D3PM [46] and ARDMs [108], extends this framework to sequences and other symbolic data [109, 110].

Recent work on Masked Diffusion Language Models (MDLM) [47] shows that a simple masked discrete diffusion process is highly competitive for language modeling. Building on this paradigm, large diffusion language models such as Dream [111], LLaDA [95], and MMaDA [112] further demonstrate that discrete diffusion can be scaled to large models and can match or even surpass strong left-to-right autoregressive baselines across a broad range of language and multimodal tasks.

Importantly, antibody sequences differ fundamentally from natural language in that they do not admit a semantically meaningful left-to-right ordering. Functional regions such as CDRs and framework residues are distributed across the sequence, and antigen binding arises from their collective paratope–epitope interactions under global structural and biophysical constraints. Moreover, long-range epistatic couplings, especially between heavy and light chains, play a critical role in determining specificity. These properties align naturally with diffusion language models, which avoid a fixed generation order and instead learn to recover globally consistent sequences through iterative denoising over randomly masked positions. As a result, diffusion language modeling provides a principled and flexible framework for learning, representing, and generating antibody sequences.

Let 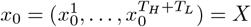 denote the concatenated paired antibody sequence defined in the previous section, where each 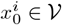 is a one-hot token over the vocabulary. The forward noising process independently corrupts each token by replacing it with the special `[MASK]` symbol:

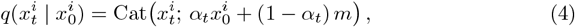

where *m* denotes the one-hot vector corresponding to `[MASK]`, and *α*_*t*_ ∈ [0, 1] is a monotonically decreasing noise schedule. This produces a partially masked paired sequence 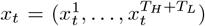. Ophiuchus-Ab is trained as denoiser to recover the original residues at masked positions, yielding the denoising objective

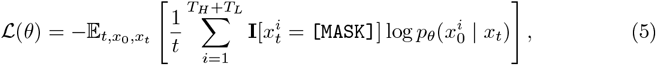

where *t* is sampled uniformly from [0, 1], *x*_0_ is the ground-truth paired antibody sequence, and **I**[*·*] denotes the indicator function.

To further improve modeling flexibility, learning efficiency, and robustness, we introduce three complementary design choices within the diffusion language modeling framework.

#### 4.2.3 Unified Training

To facilitate flexible sampling and accommodate diverse downstream design tasks, we assign independent noise levels, *t*_*h*_ and *t*_*l*_, to the heavy and light chains, respectively. This allows the model to observe and learn from asymmetric corruption patterns between the two chains, enabling it to model joint, marginal, and conditional distributions over paired antibody sequences. As a result, Ophiuchus-Ab naturally supports a wide range of applications, including CDR infilling, antibody humanization, and light-chain pairing.

During training, with a probability of 0.125, we randomly mask all tokens from either the heavy or light chain. This allows the model to handle both full antibody sequences and single-chain data, as the missing chain is masked when only one chain is available. This flexibility enables our model to tackle tasks involving singlechain sequences. Moreover, this approach facilitates the use of classifier-free guidance (CFG) [113] during inference to further improve sampling performance.

#### 4.2.4 Emphasize the modeling of the non-germline residues

To prioritize the learning of non-germline residues, particularly those within the CDR regions, we replace the standard cross-entropy loss used in traditional diffusion language models [109, 110] with focal loss [50]. Focal loss dynamically down-weights the loss for well-predicted labels, thereby focusing the model’s attention on more challenging predictions, such as the CDR regions [49]. This approach ensures that the model effectively learns the crucial non-germline mutations, which are essential for antibody understanding.

#### 4.2.5 Complementary Masking

The random masking in diffusion training means that each noised sample provides only a partial learning signal. To improve data efficiency, we adopt complementary masking [51], which ensures 100% token utilization and accelerates convergence. The core principle is to generate two antithetical training instances from a single source sequence *x*_0_. Specifically, for each source sequence *x*_0_, we generate two complementary noised sequences: a primary sequence *x*_*t*_ using a random mask, and a complementary sequence 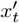 using its logical inverse. Training on both *x*_*t*_ and 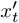 in the same batch guarantees that every token is observed in its unmasked form exactly once. This eliminates token-level sampling bias and provides a more uniform and robust learning signal at each optimization step.

#### 4.2.6 Sampling

As illustrated in Fig. 1b, we discretize the reverse diffusion process to sample from the model distribution *p*_*θ*_(*x*_0_), starting from a fully masked sequence. The total number of sampling steps is treated as a hyperparameter, providing a natural trade-off between sampling efficiency and generation quality. By default, timesteps are uniformly spaced. At each reverse step from time *t* ∈ (0, 1] to *s* ∈ [0, *t*), the current sequence *x*_*t*_ is fed into Ophiuchus-Ab to simultaneously predict all masked tokens. To obtain the next state *x*_*s*_, we then remask a fraction 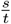 of the predicted tokens in expectation, ensuring that the reverse transition remains consistent with the forward noising process [46, 47].

In principle, the remasking operation is performed uniformly at random. However, recent work [52] has shown that diffusion language models can suffer from imbalanced learning across subproblems of varying difficulty. As a consequence, naive random unmasking during inference may over-rely on poorly trained marginals, leading to unstable or suboptimal generations. To mitigate this issue, we adopt adaptive sampling strategies that prioritize the unmasking of low-confidence predictions, resulting in more reliable and stable generation.

This sampling framework naturally generalizes to a wide range of antibody design tasks. Unconditional sequence generation is achieved by initializing the process from a fully masked sequence, while conditional tasks such as CDR or other region infilling are realized by masking only the target regions and keeping the remaining tokens fixed. The reverse diffusion process then fills in the masked positions in a globally consistent manner, without requiring task-specific model modifications or specialized decoding procedures.

### 4.3 Implementation details

The models were trained on 8 NVIDIA A100 GPUs with mixed-precision training enabled. The global batch size was set to 128, corresponding to a per-GPU batch size of 16. Optimization was performed using AdamW with a learning rate of 4 × 10^−5^, *β*_1_ = 0.9, *β*_2_ = 0.98, and a weight decay of 0.01. We employed a polynomial learningrate scheduler with 2k warm-up steps, and trained the model for a total of 100k optimization steps.

## Declarations

### Data availability

We pre-trained Ophiuchus-Ab on paired antibody sequences from the OAS database [21] using custom training splits, and evaluated it on the following datasets:

- SAbDab [20]
- GDPa1 [83]
- Binding affinity [87]
- Antibody specificity [93]

### Code availability

The inference source code of this work is available at https://github.com/Ophiuchus-Team/Ophiuchus-Ab.

